# Paclitaxel-induced peripheral neuropathy is caused by epidermal ROS and mitochondrial damage through conserved MMP-13 activation

**DOI:** 10.1101/743419

**Authors:** Anthony M. Cirrincione, Adriana D. Pellegrini, Jessica R. Dominy, Marisa E. Benjamin, Irina Utkina-Sosunova, Francesco Lotti, Stanislava Jergova, Jacqueline Sagen, Sandra Rieger

## Abstract

Paclitaxel is a chemotherapeutic agent that causes peripheral neuropathy (PIPN) as a side effect of cancer treatment. Severely affected patients need to terminate chemotherapy, diminishing their chance of survival. The underlying causes of PIPN are unclear, but epidermal, unmyelinated axons have been shown to be the first to degenerate. We previously utilized a zebrafish *in vivo* model to show that the epidermal matrix-metalloproteinase 13 (MMP-13) induces degeneration of unmyelinated axons, whereas pharmacological inhibition of MMP-13 prevented axon degeneration. The precise functions by which MMP-13 is regulated and affects axons, however, remained elusive. In this study, we assessed mitochondrial damage and reactive oxygen species (ROS) formation as possible inducers of MMP-13, and we analyzed MMP-13-dependent damage. We show that the small ROS, H_2_O_2_, is increased in keratinocytes following treatment with paclitaxel. Epidermal mitochondrial damage appears to be a source of ROS leading to cytoplasmic H_2_O_2_ elevation, upregulation of MMP-13, and increased matrix degradation. Intriguingly, although axonal mitochondria also show aberrant morphologies and are vacuolized, as shown in other neuropathies, these axonal mitochondria do not produce increased H_2_O_2_ levels. We suggest that mitochondrial vacuolization occurs independently of axonal damage given that MMP-13 inhibition prevents axon degeneration, though vacuoles persist. We further show that MMP-13 dysregulation also underlies PIPN in rodent paclitaxel models, and that this function appears to be DRG neuron-extrinsic. These findings suggest that vacuolization is not a cause of paclitaxel-induced neuropathy, and that epidermal MMP-13 is a strong candidate for therapeutic interventions in cancer patients with neuropathy.

## Introduction

Approximately 60-70% of cancer patients treated with paclitaxel develop peripheral neuropathy (1). This condition affects predominantly somatosensory axons in the palms and soles of hands and feet, leading to pain, tingling, temperature sensitivity, and numbness (2, 3). Unmyelinated terminal arbors of somatosensory axons innervating the epidermis have been suggested to be the first to degenerate (4), though the mechanisms leading to their degeneration has remained elusive. We previously showed the conservation of this process in zebrafish, and we mechanistically demonstrated that axon degeneration is mediated by overactivation of the matrix-metalloproteinase, MMP-13 (5). However, the precise mechanism of MMP-13 regulation and the role of MMP-13 in the induction of axon degeneration remained unclear. Scanning electron microscopy indicated that paclitaxel treatment promotes epidermal cell abrasions and adhesion defects, suggesting that increased matrix degradation in the epidermis by MMP-13 may be a cause of axon degeneration whereby axons detach from the matrix due to continuous degradation processes and mechanical stress on the epidermis.

MMP-13 is a member of the matrix-metalloproteinase family of matrix-degrading enzymes. This family has been shown to be regulated by reactive oxygen species (ROS), such as hydrogen peroxide (H_2_O_2_). For instance, treatment of prostate cancer cells with H_2_O_2_ elevated *mmp3* expression levels through inhibiting the MMP-3 suppressor, Thrombospondin 2, in a microRNA-dependent manner (6). MMPs can be particularly regulated by mitochondrial ROS (mtROS). For instance, MCF-7 breast cancer cells treated with the mtROS inducer, rotenone, showed increased ROS production and *mmp2* expression. This effect was dependent upon manganese superoxide dismutase (7). The mitochondrial ROS-dependent regulation of MMPs is especially interesting given that paclitaxel treatment directly targets mitochondria, such as in cancer cells (8), and also upregulates MMP-13 in basal keratinocytes in our zebrafish model (5). Since paclitaxel shows strong efficacy in the treatment of carcinomas, an epithelial-derived cancer cell type, this chemotherapeutic agent could similarly induce mitochondrial dysfunction in basal epidermal keratinocytes, leading to MMP-13 upregulation and axon degeneration. Here we assess this idea and analyze how MMP-13 contributes to the degeneration of unmyelinated sensory axons innervating the epidermis.

## Materials and Methods

### Animals

Zebrafish (Nacre and AB, AB/TL, or Tu/TL strains) were raised and bred according to NIH guidelines and handled in strict accordance with good animal practices as approved by the appropriate IACUC committees. Zebrafish eggs were collected in a strainer and rinsed with deionized water, followed by transfer into Petri dishes filled with Instant Ocean medium +Methylene Blue. The eggs were cleaned after 24hr and placed into Ringers solution (Zebrafish Book Protocols) with or without Phenylthiourea (Sigma Aldrich) if pigmented. The embryos were kept at 28.5°C in a 14:10hr light cycle incubator until experimentally used.

Rat and mouse behavioral studies were completed at the University of New England Behavioral Core Facility and at the University of Miami, according to IACUC regulations at each institution. Housing and maintenance conditions met NIH/ OLAW standards. Seven-week old Sprague-Dawley rats were obtained from Envigo, whereas C57BL/6J mice were obtained from The Jackson Laboratory. Upon arrival, rodents were acclimated for 3-5 days. Rats were housed in groups of two, whereas mice were housed in groups of 5. The animals were subsequently handled for 3 consecutive days to accustom them to the testing conditions. Following this initial period, the animals were habituated to testing chambers and baseline behavior measurements were obtained for von Frey (tactile), Hargreaves (thermal) and cold plate or acetone assays, as described below.

### Plasmids and transgenic lines

UAS-mito-dsRed fish (gift from Dr. Alvaro Sagasti (UCLA)) were injected with *krt4*:Gal4VP16 to label skin mitochondria and CREST3:Gal4VP16_14×UAS-GFP to label sensory neurons. *krt4*:HyPer-cyto was previously published previously (1). *tp63*:HyPer was generated by amplifying HyPer-cyto from the original Evrogen plasmid with 5’ BamHI and 3’ NotI sites via PCR. The HyPer amplicon was inserted into *tp63*:AcGFP (gift of Dr. Gromoslav Smolen) after AcGFP was removed via BamH/NotI digestion. CREST3:Gal4VP16_14×-HyPer-cyto was cloned using the Gateway system (Life Technologies). pDONR vectors for CREST3 and Gal4VP16-14×UAS were a gift from Dr. Alvaro Sagasti (UCLA). HyPer-cyto was amplified via PCR with attB primers to insert as 3’ element into the pDEST vector together with CREST3 and Gal4VP16-14×UAS using recombination. Transgenic *isl2b*:GFP fish (2) were used for axon degeneration studies.

### Drug administrations

#### Zebrafish treatments

Starting at 2dpf, zebrafish were placed individually into each well of 24-well plates. The control groups were typically incubated in 0.09% DMSO/Ringers solution to match the DMSO concentration in the final paclitaxel solution. Pre-made paclitaxel solution was purchased from Millipore as a 25mM stock solution in DMSO. DB04760 (MMP-13 inhibitor, PubChem CID: 5289110) (sc-205756, Santa Cruz Biotechnology) and CL-82198 (Tocris, USA) were purchased as lyophilized powder and diluted with DMSO as 10mM stock solutions that were stored at −20°C. Paclitaxel was diluted to a final concentration of 22-30μM (as indicated in the text) and DB04760 and CL-82198 were diluted 1:1000 to 10μM solutions in Ringers medium prior to each use. Rotenone and paraquat were purchased from Sigma-Aldrich and stored as 10mM stock solutions in DMSO until use. Rotenone was diluted to 0.1μM and paraquat to 10μM in Ringers. N-acetylcysteine was used at 1.5mg/L final concentration. If necessary, the solutions were exchanged every 2 days into fresh ones, as indicated in the text. All animals in treatment solutions were kept in the dark in the incubator until the beginning of the analysis, to avoid photo-toxicity and reduce degradation. Light exposure was minimized for animals injected with HyPer during the imaging procedure to avoid photobleaching and oxidation.

#### Rodent treatments

Animals were divided into four groups: vehicle (Cremophor-EL/EtOH 50/50 diluted into 0.9% NaCl), paclitaxel/vehicle, vehicle+CL-82198 or paclitaxel+CL-82198. Paclitaxel was injected at 2mg/kg on four alternating days (D1, D3, D5 and D7) into rats and at 4mg/kg into mice for the same duration. Cremophor-EL is a derivative of castor oil and ethylene oxide that is used clinically for paclitaxel injections. Paclitaxel/Cremophor was prepared as a stock concentration of 8mg/mL and injected intraperitoneally (i.p.) using an injection volume of 1ml/kg. CL-82198 was diluted in 0.05% DMSO/PBS and injected daily for 7 days at a concentration of 1, 5, or 10mg/kg. For topical application of CL-82198, rats were injected with vehicle or paclitaxel as stated above. The ipsilateral hind paw relative to the injection site was treated with 10μM CL-82198 in 0.01% DMSO once a day during the injection period (for seven days).

### Western blot

Pools of 10 zebrafish at 4dpf were collected after two days of treatment with compounds indicated in the text. The larvae were pipetted into 2ml microcentrifuge tubes, depleted of any medium, and frozen in liquid nitrogen. The larvae were stored at −80°C until use (~1-4 weeks). For western blot analysis, frozen zebrafish were placed on ice and incubated in 200μL 0.5× RIPA lysis buffer (MilliPore) containing 1× cOmplete™ Protease Inhibitor Cocktail (Roche), followed by 10sec homogenization using a ProScientific Bio-Gen Pro200 homogenizer.

Samples were combined with 4× Laemmli sample buffer (Invitrogen) and reducing agent (Invitrogen) before separation on SDS-PAGE 4-12% Bis-Tris gels (Invitrogen). Samples were transferred to a 0.45 μM PVDF membrane (Invitrogen) using a Trans-Blot Semi-Dry apparatus (Bio-Rad) according to the manufacturer’s protocol. The membranes were incubated for 1hr in blocking buffer (Odyssey) at room temperature before incubation with primary MMP-13 rabbit polyclonal antibody (ProteinTech, 1:333), and monoclonal beta-actin antibody (Abcam ab8226, 1:2000) overnight at 4°C rocking. After three 5-minute washes in TBSTween (0.1%) at room temperature, the membranes were incubated in secondary fluorescent antibodies (anti-rabbit IgG 1:5000 IRDye 800RD; anti-mouse IgG 1:5000 IRDye 680CW) in blocking buffer + 0.01% SDS and 0.2% Tween-20 (according to Odyssey protocol) at room temperature for 1 hour on a shaker. Blots were washed three times for 5 minutes in TBS-Tween (0.1%) and two times for 5 minutes in TBS before imaging on an IR imaging system (Li-Cor).

### Zebrafish transmission electron microscopy studies

Larval zebrafish were fixed in 2% Glutaraldehyde (Electron Microscopy Sciences, Cat. No. 16120)/ 2% PFA/ PBS (Fisher Scientific, Cat. No. NC0603972) overnight at 4°C and washed 3 times in PBS. The samples were post-fixed in 2% aqueous osmium tetroxide (Sigma-Aldrich, Cat. No. 201030), rinsed in PBS, and then dehydrated with ethyl alcohol (Sigma-Aldrich, Cat. No. E7023). Samples were subsequently infiltrated with Embed 812 resin (Electron Microscopy Sciences, Hatfield, PA), and the fish were embedded such that the caudal fin could later be sectioned in transverse orientation (axially) starting from the caudal end and moving anteriorly. Following orientation of one larva in each mold, the mold was slowly filled with resin and polymerized in a 60°C oven for 24hr. 90nM sections were cut from each block on a Leica UC6 ultra microtome and collected on 300 mesh copper grids (Sigma-Aldrich, Cat. No. G4901-1VL). After staining with 1% aqueous uranyl acetate/Reynold’s lead citrate, grids were viewed on a JEOL JEM 1230 transmission electron microscope and images were collected with an AMT 2K digital camera.

### Rat tissue harvesting and immunostaining

Male Sprague-Dawley rats were euthanized with CO_2_ and intracardially perfused with saline, followed by 4% paraformaldehyde. Spinal cords were dissected, post fixed in 4% paraformaldehyde overnight at 4°C, and transferred into 30% sucrose solution for cryoprotection for 48hr. Lumbar spinal cord tissue was embedded in the cryoprotective media, sectioned on the cryostat with 40μm thickness, and collected into 1×PBS in series. Sections of the lumbar L3-L5 spinal cord were processed for immunostaining to visualize CGRP fibers in the dorsal horns. Briefly, tissue sections (at least 10 per animal from 5 animals) were incubated in 5% normal goat serum (NGS) in PBS with 0.05% Triton X-100 (PBSTx) for 2hr followed by overnight incubation with CGRP (1:1000, Millipore) at 4°C. Sections were washed 4×5min with PBS and incubated in secondary antibody (1:250, Alexa flour-633, ThermoFisher) in 5%NGS and PBSTx for 2hr, washed in PSB, incubated with DAPI for nuclear staining (1:1000, Invitrogen), washed 4×5min with PBS, mounted on slides, air-dried overnight and cover-slipped with Vectashield media (Vector labs).

### Confocal imaging

Live larval zebrafish were mounted according to (3) and imaged on an Olympus FV1000 confocal microscope, or a Zeiss Apotome microscope with a 20× air objective. Sections were recorded in 3-10μm steps. Images were processed in Imaris (Bitplane) for quantifications. ZEN software was used for HyPer measurements and division image display. Fiji was used to measure mitochondria in the TEM images. Prism (GraphPad) was used to generate graphs and perform statistical analyses. For publication, sections were projected into single planes using the Maximum Intensity Projection function, followed by image processing in Photoshop. Graphics were generated in Microsoft PowerPoint and Adobe Illustrator.

For imaging of rat sections, slides were examined under a fluorescent microscope (Zeiss Axiovert 200m for light and fluorescence microscopy) at 20× magnification and digitalized by attached camera. Images were processed by ImageJ software. Background intensity of each image was manually adjusted to reduce variability. Images were then converted into black and white mode and optical density of the dorsal horn (Lamina LI-LV) was measured by the software. This method was chosen to evaluate not only the changes in the superficial dorsal horn but also to reflect possible sprouting into deeper dorsal horn laminae. Representative images were taken with confocal microscope Olympus Fluoview 100 (Olympus Spectral Confocal Microscope Fluoview 1000 equipped with multiple lasers covering full spectrum from 400nm – 680nm).

### Adult mouse DRG cultures

*Dissection and Harvest of Adult Mouse DRG Neurons*: After anesthetizing 8-week adult mice with 0.5% isoflurane, mice were decapitated to remove the dorsal skin and spinal column. The removed spinal column was washed 2-3 times with 1× PBS and pinned with the ventral side up to a Petri dish filled with chilled PBS. Under a dissecting scope, the muscles were carefully removed, and a laminectomy performed. The spinal cord was removed from the ribs by cutting the roots and leaving DRGs in the spinal column. This was followed by slicing the dorsal side of the spinal column longitudinally in two halves and reattaching them with pins. DRGs were collected and maintained in 3cm Petri dishes with HBSS (w/o calcium and magnesium) on ice. Under a dissecting scope, the roots and meninges were trimmed from the DRGs. DRGs from cervical, thoracic and lumbar anatomical parts were harvested. *Digestion and Dissociation*: After collecting the dissected DRGs, they were transferred into a microcentrifuge tube containing 1mL collagenase A (2.5mg/ml) in MEM medium, and incubated at 37°C for 40 min for digestion. The collagenase A solution was replaced with 1mL of 1×TrypLE Express (trypsin-containing) solution and DRGs were incubated at 37°C for 10 min. After the trypsinization step was stopped with 10% FBS, DRGs were spun at low speed to remove the trypsin. DRGs were transferred into a 15ml tube containing 2mL MEM medium supplemented with 100U/ml DNAse I and gently pipetted up and down 10 times for trituration, using a 1mL graduated pipette tip. After trituration, the non-dissociated tissues were allowed to settle down to the bottom of the tube and the cell suspension was transferred to a new 15ml sterile tube. The trituration step was repeated until most of the tissue was dissociated. The cell suspension was combined to a total volume of 5mL. The obtained cell suspension contained both neurons and non-neuronal cells. Neuronal cells were therefore further purified via a layer of 4% BSA by centrifugation, followed by resuspension of the pellet in 0.5mL MEM, followed by quantification of the neurons with a hemocytometer. Neuronal cells were plated at a desired density of ~50 DRGs (number of neurons ~7 ×10□) in MEM medium supplemented with 1× MEM vitamin supplement, 1× B27, 1% penicillin/streptomycin and 20μM 5-fluorodeoxyuridine/uridine. Usually, neuronal cells were plated in 96-well plates coated with 100μg/ml polyornitine and 10μg/ml laminin.

### Behavioral studies

Cold response, Hargreaves (thermal response) and von Frey filament testing was conducted on the following days: pre-injection, day 2, 4, 7, 9, 11, 14, 16 and 23. Animals were sacrificed on day 23.

#### Cold-plate test (USDA Category C)

Rats were acclimated to the room for 30 minutes. The cold plate (Ugo Basile) was used and set to a consistent temperature of 5°C. Distilled water was added onto the plate, making the surface wet prior to testing. One rat was tested at a time and once the rat was on the plate the timer was started.

The time taken in seconds for each rat to respond was recorded. A response was recorded as removal of the hind paw from the cold plate (shaking, licking, lifting or jumping). Only the hind paw responses were considered, as front paw movements were mostly attributed to grooming behavior of the mice. The test was stopped after 300 seconds if no response was observed. Cold latencies (seconds) were recorded for each rat and group means and the Standard Error of the Mean (SEM) was calculated and graphed.

#### Acetone test to assess cold allodynia (Figure 4H)

Sensitivity to a non-noxious cooling stimulus was evaluated using acetone. One hundred μl of acetone was dropped onto the lateral margin on the hind paw from a blunted 22g needle attached to a syringe. In normal, intact rats, acetone does not evoke a withdrawal response. Acetone was applied to the hind paw 5 times, with about 1-2 min between applications. The total number of positive responses out of five was converted to a percent response frequency (4). The response was marked as positive only when a supraspinally driven reaction occurred in addition the paw withdrawal, such as a head turning towards the stimulus, paw licking, or shaking.

#### Thermal hyperalgesia (USDA Category C)

For assessment of thermal withdrawal latencies, animals were acclimatized for ~30 min to Plexiglas holding chambers that rest on a glass surface maintained at room temperature. The surface was cleaned of urine and feces prior to the assessments. Thermal nociceptive thresholds were determined similar to the methods described by Hargreaves et al. (5). Briefly, a radiant heat source (Ugo Basile) was focused through the glass surface onto the plantar surface of the hind paw. Upon paw withdrawal, the heat stimulus was automatically deactivated and the latency to withdraw was recorded to the nearest 0.1 seconds. Each latency score is an average of two trials separated by at least 5 minutes. Both hind paws were typically tested. The intensity of the light stimulus was set for a baseline latency of approximately 20-25 seconds. The test was terminated to prevent tissue damage in the absence of a response within a predetermined maximum latency (30 seconds). Escape attempts (jumping) were not typically observed except after multiple hot-plate exposures and/ or higher hot-plate temperatures. We used as criteria only hind paw responses since forepaw licking and lifting are components of normal grooming behavior. Each animal was tested only once since repeated testing in this assay leads to profound latency changes (e.g.(6) (7) (5). We also employed a strict baseline, and cut-off times (e.g., 30 seconds) to prevent tissue damage in animals not responding to the stimulus.

#### Von Frey Filament Testing of Mechanical Sensitivity (USDA Category C)

Animals were placed in a suspended plastic chamber with a wire mesh platform and allowed to habituate. Tactile hypersensitivity was determined by measuring paw withdrawal thresholds in response to probing the plantar surface of the left hind paw with a series of 5 calibrated von Frey filaments (4.31, 4.56, 4.74, 4.93, and 5.18). The von Frey filaments were pressed into the soft spot on the foot and pressed upward until the filament bent in half for a duration of 5 seconds. If the rat expressed no response (absent licking or withdrawal of the hind paw), the response was marked as “O” and the next highest filament was tested. If a response occurred, the test was terminated, and the response marked as “X”. Tactile thresholds (g) were calculated for the “XO” pattern of each rat. Measurements were taken once daily on indicated days before the initial injection (baseline) and at later days following paclitaxel and CL-82198 injections. Withdrawal thresholds were determined by sequentially increasing and decreasing the stimulus intensity (“up and down” method). The data was analyzed using a Dixon nonparametric test and expressed as the paw withdrawal threshold in gram force values (8). The group means and standard error of the mean (SEM) was calculated and graphed.

### Statistical analyses

Statistical comparisons were made using Prism7 (GraphPad) using Student’s t-test and one-way ANOVA at an alpha=0.05 (95% confidence interval) and Tuckey’s or Bonferroni’s multiple comparisons post-test, as indicated. Significance is denoted with asterisks: *p<0.05, **p<0.01, ***p<0.001, ****p<0.0001. The standard deviation (S.D.) or error of the mean (S.E.M.) is shown, reflecting comparisons of two groups in a single experiment or multiple biological replicates, respectively.

## Results

A prevalent model for paclitaxel neurotoxicity posits that paclitaxel causes axon degeneration by intra-axonal mitochondrial damage and ROS formation (9–11), which parallels findings in cancer cells where paclitaxel treatment *in vitro* induces mitochondrial damage and ROS, ultimately inducing cancer cell apoptosis (8). However, it remains unclear whether the observed mitochondrial damage in axons is a cause of axon degeneration or the consequence of degradation processes induced during axon degeneration (**Figure 1A**). *In vivo* analyses will be useful to dissect this question in more detail using fluorescent genetic H_2_O_2_ sensors and mitochondrial markers.

**Figure 1.**
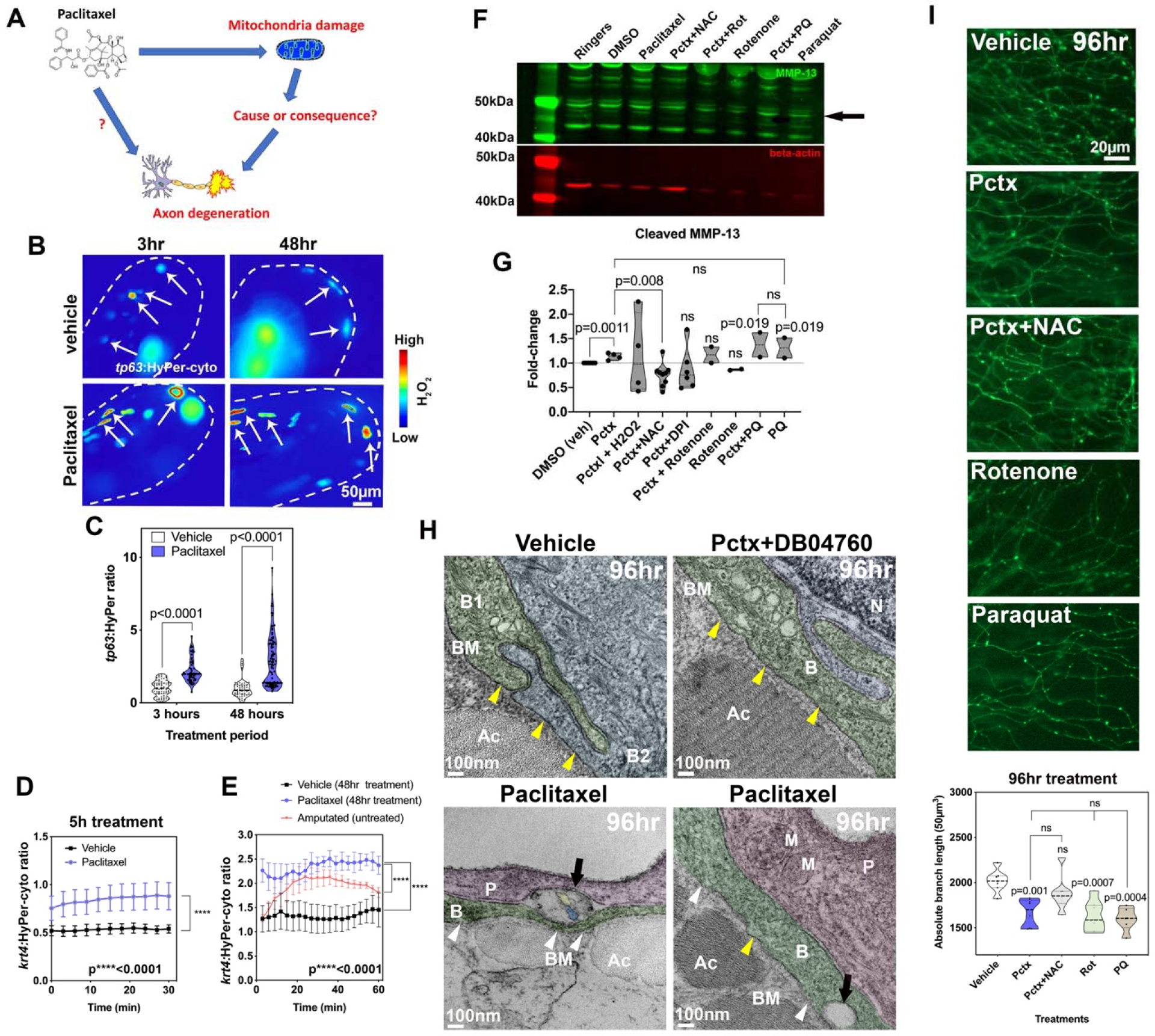
Mitochondrial ROS contribute to MMP-13 expression and axon degeneration. **(A)** Is mitochondrial damage involved in paclitaxel-induced axon degeneration? **(B)** Ratiometric images showing HyPer oxidation (arrows) in the caudal fin of larval zebrafish (dashed lines) after 3 and 48hr of treatment (2 and 4dpf, respectively) with either 0.09%DMSO vehicle or 23μM paclitaxel. Keratinocytes are mosaically labeled following transient injection of *tp63*:HyPer into 1-cell stage embryos. High oxidation is represented in red and low oxidation in blue. **(C)** Quantification reveals increased oxidation after short and long-term treatment with paclitaxel, n=10 animals. **(D, E)** 30-60 min quantifications of HyPer oxidation in epidermal cells expressing *krt4*:HyPer following treatment for 5 (**D**) and 48hr (**E**), n=3-6 animals, SEM. Paclitaxel treatment significantly elevates H_2_O_2_ levels. Comparison with H_2_O_2_ induced by fin amputation shows continuous versus transient H_2_O_2_ elevation. **(F, G)** Western blot (black arrow indicates cleaved MMP-13, 48kDa) (F) and quantitative analysis (G) shows increased MMP-13 expression following treatment with paclitaxel and the mitochondrial ROS inducer paraquat but not rotenone. Antioxidants (NAC and DPI) reduce MMP-13 levels induced by paclitaxel treatment, n=2 biological replicates with 5-10 animals each treatment. **(H)** TEM of (6dpf) zebrafish keratinocytes following 96hr treatment with either 0.09% DMSO vehicle, 23μM paclitaxel+DB04760 (MMP-13 inhibitor) or paclitaxel alone shows an intact basement membrane (yellow arrowheads) in vehicle and paclitaxel+DB04760 treated fish, whereas the basement membrane is discontinuous (white arrowheads) and large gaps (black arrows) appear where unmyelinated sensory axons normally reside. **(I)** Epidermal unmyelinated sensory axons in the distal tail fin following 96hr treatment with vehicle, paclitaxel, paclitaxel+NAC, rotenone, or paraquat. Quantification of treatments shows paclitaxel, rotenone, and paraquat induce axon degeneration whereas NAC co-administration prevents degeneration, n=5-7 animals, SEM. *Abbreviations: NAC*=*N-acetylcysteine*, *Pctx*=*paclitaxel*, *Rot*=*rotenone*, *PQ*=*paraquat*, *B*=*basal keratinocyte*, *BM*=*basement membrane*, *Ac*=*actinotrichia*, *N*=*nucleus*, *P*=*periderm*, *M*=*mitochondrion*, *dpf*=*days post fertilization*

We first wanted to analyze whether paclitaxel causes H_2_O_2_ formation in epidermal keratinocytes and whether H_2_O_2_ regulates MMP-13 expression. For this, we specifically expressed the H_2_O_2_ sensor, HyPer (12), in keratinocytes under the *krt4* and *tp63* promoters (5). The *krt4* promoter drives expression in both epidermal layers and is later restricted to differentiated keratinocytes of the surface periderm layer. The *tp63* promoter is restricted to basal epidermal keratinocytes with expression starting around 24hpf when the basal layer forms. HyPer oxidation was measured and represented as the ratio of oxidized to non-oxidized HyPer (**Figure 1B-D**). HyPer oxidation was significantly increased in *tp63* basal keratinocytes of zebrafish treated with paclitaxel over short (3hr) and long-term (2-day) periods (**Figure 1B, C**, **S1**). A similar elevation was observed when HyPer was expressed for 5hr and 48hrs under the *krt4* promoter (**Figure 1D, E**). This suggests that paclitaxel elevates H_2_O_2_ levels in both keratinocyte layers.

Previous studies suggested that wounding such as by fin amputation induces H_2_O_2_ production in the epidermis (13), and we showed that this process promotes axon regeneration (14). We, therefore, wondered why oxidation in this context is not toxic but pro-regenerative. By comparing amputation induced H_2_O_2_ levels to those induced by paclitaxel, we noticed that paclitaxel treatment led to continuous H_2_O_2_ production at steady state in comparison to a transient rise of H_2_O_2_ during the initial ~20min after amputation followed by declining levels thereafter (**Figure 1E**). Thus, it appears that axons and skin cells can cope with some exposure to H_2_O_2_, such as during an injury response, likely due to rapid activation of antioxidant complexes after the initial H_2_O_2_ production. However, long-term exposure has an opposing effect.

We next wanted to know whether H_2_O_2_ regulates MMP-13 expression in the context of paclitaxel treatment using western blot analyses. For this, we treated zebrafish either with 0.09% DMSO vehicle (matching the percentage of DMSO contained in the paclitaxel), 23μM paclitaxel plus or minus the antioxidant 1.5mg/L N-acetylcysteine (NAC), and the mitochondrial mtROS inducers rotenone (0.1μM) and paraquat (10μM) for two days (**Figure 1F, G**). The drug doses (except DMSO) reflect the maximally tolerated dose. Compared to vehicle-treated fish, paclitaxel induced the expression of MMP-13 (as previously shown (5)) and this upregulation was reduced below control levels in the presence of NAC. Rotenone and paraquat had the opposite effects. However, rotenone treatment without paclitaxel did not appear to induce MMP-13 expression, which might be a dose-dependent effect. It is also possible that because rotenone induces apoptosis, other pathways in the absence of paclitaxel contributed to the inhibition of MMP-13 expression. Paraquat with and without paclitaxel appeared to increase MMP-13 expression at comparable levels to paclitaxel, suggesting that mtROS might be the sole source of ROS contributing to its expression, whereas cytoplasmic NADPH oxidases are not involved.

To determine how MMP-13 activity affects skin morphology and axon degeneration, we performed transmission electron microscopy (TEM) in zebrafish treated for 96hr (until 6 days post fertilization, dpf) with paclitaxel with and without the MMP-13 inhibitor, DB04760. Treatment with paclitaxel for this period results in prominent axon degeneration (5). This showed paclitaxel-induced basement membrane defects, evidenced by a discontinuous, electron-dense barrier (yellow arrowheads) that separates epidermal basal cells from the underlying dermis (15) (**Figure 1H**). Overall, the tissue appeared less electron-dense, and the keratinocytes in paclitaxel-treated zebrafish appeared loosely attached, as opposed to their tight connections seen in control animals. None of these defects were present when paclitaxel and DB04760 were co-administered. These findings indicate matrix defects as a likely cause of axon degeneration. A pressing question in the field is why axons first begin to degenerate in the hands and feet. Our data suggests the possibility that a combination MMP-13 dependent matrix degradation and increased mechanical stress in the palms and soles might lead to the detachment of axons from the matrix and ultimately causes axonal degeneration.

To determine whether mtROS also cause axon degeneration, consistent with mtROS-dependent MMP-13 upregulation, we quantified intraepidermal branch length following treatment for two days with either vehicle, paclitaxel, paclitaxel+NAC, paclitaxel+rotenone, or paclitaxel+paraquat. We found significant axon degeneration upon paclitaxel treatment with and without rotenone and paraquat, compared with vehicle controls and paclitaxel+NAC treatment, suggesting that paclitaxel induces axon degeneration through mtROS in an MMP-13-dependent manner (**Figure 1I**).

We next wanted to assess whether epidermis-specific mitochondrial damage specifically was the underlying source of mtROS. TEM analysis of mitochondria in basal keratinocytes showed that the cristae and outer and inner membranes appeared less distinct after paclitaxel treatment compared with controls (**Figure 2A, B**, **S2**, **S3**). We also captured one fusion event of two mitochondria following paclitaxel treatment, which we never observed in the controls (**Figure S3**). Quantification of mitochondria diameter (**Figure 2C**) revealed several effects: vehicle-treated mitochondria appeared smaller at 6dpf compared with 2dpa, which could relate to long-term effects of DMSO treatment, or be the result of ongoing developmental changes in the animals, as has been described by others (16). Interestingly, 3hr treatment with paclitaxel led to an acute shrinkage of mitochondria compared with vehicle controls. This effect could not be prevented by co-administration of MMP-13 inhibitors, DB04760, and CL-82198. In contrast, prolonged treatment with paclitaxel led to mitochondrial swelling, whereas co-administration with either of the two MMP-13 inhibitors alleviated the swelling and restored mitochondrial morphology (**Figure S4**). It is possible that the epidermis remained intact upon MMP-13 inhibition, which also prevented perturbations of mitochondrial function. Thus, mitochondria damage upon long-term exposure to paclitaxel appears to be secondary, while early defects could be a direct effect of paclitaxel.

**Figure 2.**
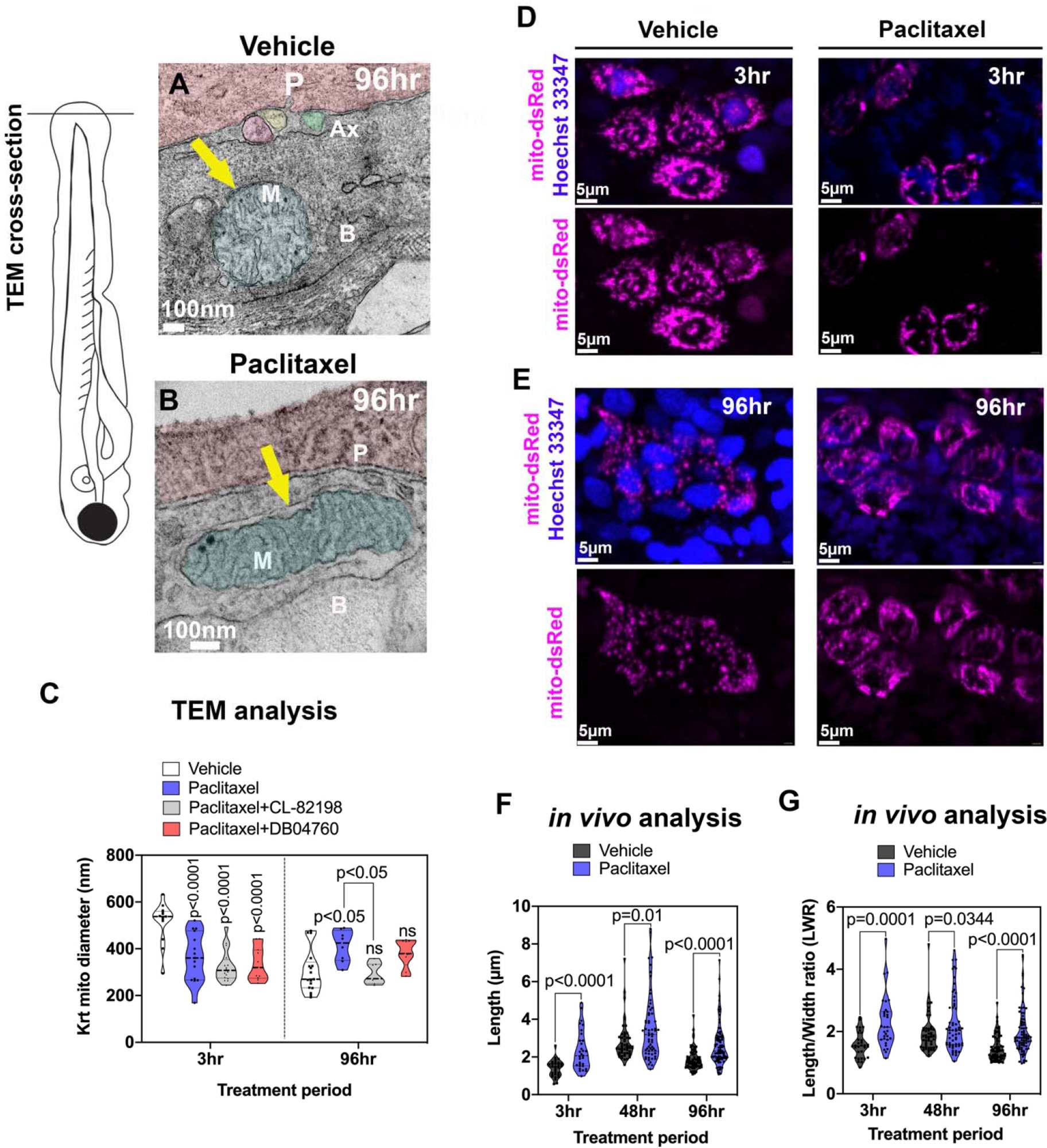
Keratinocyte mitochondria are damaged by paclitaxel treatment. **(A)** TEM analysis of zebrafish larvae treated for 96hr with either 0.09% DMSO vehicle or 23μM paclitaxel (2-6dpf). A mitochondrion (M) is shown with outer and inner membrane (white arrow) and clearly visible cristae following vehicle treatment. Three axons (Ax) are embedded between the periderm and basal keratinocyte layer. **(B)** The outer membrane of a mitochondrion following paclitaxel treatment is less apparent, and the space between inner and outer membrane has increased (yellow arrow). The cristae are less electron-dense, and large dark puncta are visible. **(C)** Quantification shows decreased mitochondria diameters upon 3hr paclitaxel treatment with and without MMP-13 inhibitors, CL-82198 or D04760, compared with controls. 96hr treatment shows increased mitochondria diameters in paclitaxel but not MMP-13 inhibitor treated animals, n=3 animals. **(D, E)** *In vivo* imaging of mitochondria in keratinocytes labeled with mito-dsRed and nuclei labeled with live Hoechst 33347 stain following treatment for 3hr and 96hr with either vehicle (**D**) or paclitaxel (**E**). Short and long-term treatment with paclitaxel promotes mitochondria filamentation. **(F, G)** Quantification after 3hr, 48hr and 96hr treatment shows increased length (**F**) and length/width ratios (**G**) with paclitaxel compared to controls, n=5-8 animals. *Abbreviations: B*=*basal keratinocyte*, *P*=*periderm*, *M*=*mitochondrion*, *Ax*=*axon*

To further analyze mitochondrial morphology, we tagged keratinocyte-specific mitochondria *in vivo* with mito-dsRed. Measurements of mitochondria length reveled that paclitaxel treatment induced mitochondrial filamentation when analyzed after 3hr, and this effect persisted even after four days of treatment with paclitaxel (**Figure 2D-G**, **S5**). This data indicates that paclitaxel affects mitochondrial fusion or fission in keratinocytes.

Previous studies investigated mitochondria in DRG axons of rodents treated with paclitaxel, which suggested mitochondrial swelling and vacuolization. These changes were suggested to underlie PIPN (9, 17, 18). To determine if such morphological changes are conserved in zebrafish treated with paclitaxel, we used *in vivo* imaging and TEM analysis (**Figure 3**). Tagging axons with mito-dsRed revealed significant morphological differences, similar to those observed in keratinocytes, namely increased mitochondria diameters suggestive of swelling and enhanced filamentation (**Figure 3A-I**). Intriguingly, despite the increased length of mitochondria, we counted more mitochondria per axon segment in paclitaxel-treated animals compared to controls. These results indicate a possible increase in fusion and fission events to maintain mitochondrial health.

**Figure 3.**
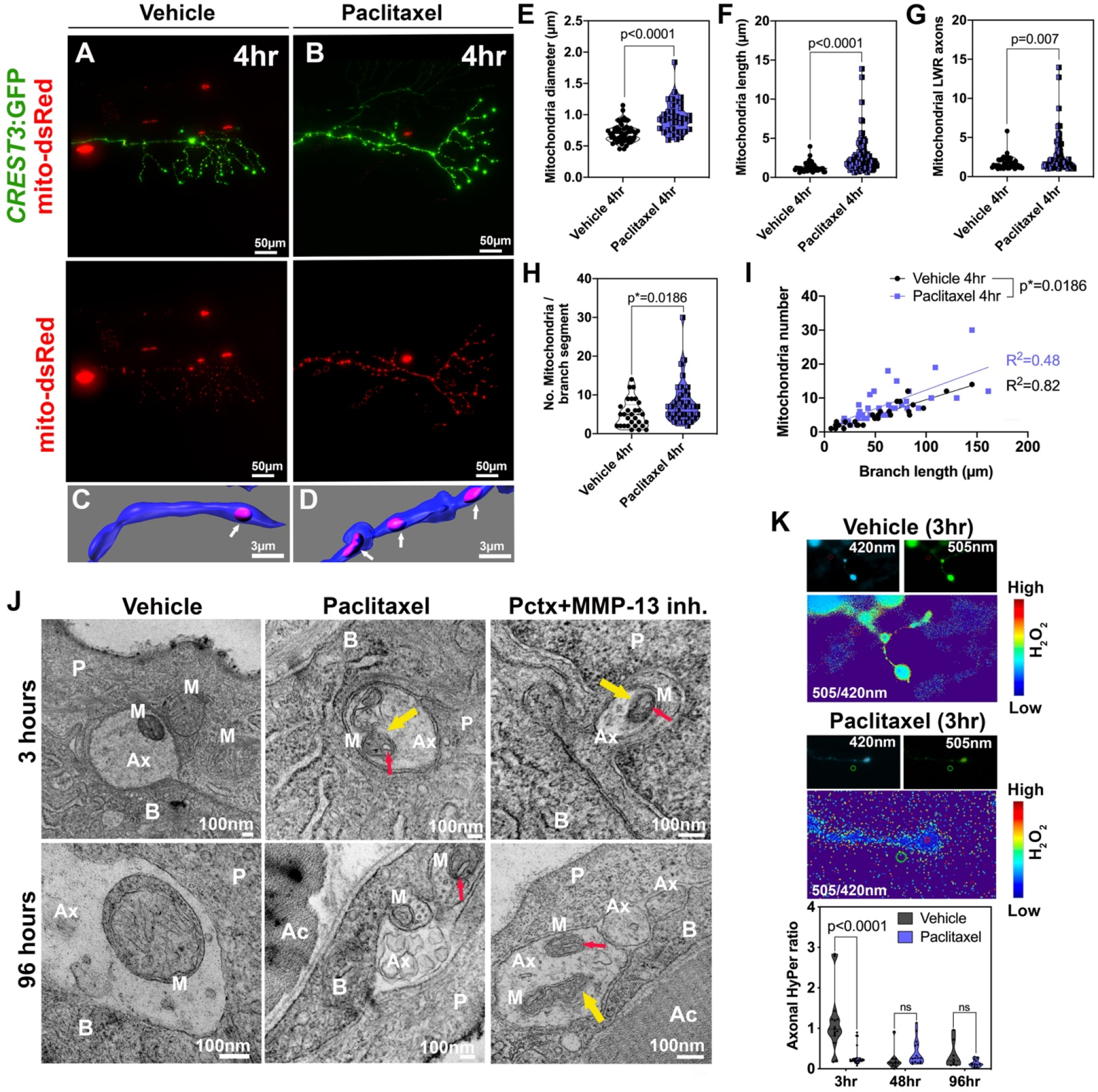
Axonal mitochondria are vacuolized following paclitaxel treatment but ROS/H_2_O_2_ levels are not elevated. **(A, B)** DsRed-labeled mitochondria in unmyelinated axons of somatosensory neurons labeled with CREST3:GFP following 4hr treatment with vehicle (**A**) or paclitaxel (**B**). **(C, D)** Respective 3D reconstructions of axon branch segment (blue) in the epidermis containing mitochondria (pink, arrows). **(E-I)** Quantification of axonal mitochondria shows increased diameter (**E**), length (**F**), length-width ratio (**G**) and number of mitochondria per branch segment (**H, I**), n=5 animals. **(J)** TEM analysis of axonal mitochondria (M) following treatment with 0.09% DMSO vehicle, paclitaxel with or without MMP-13 inhibitor (Cl-82198 top and DB04760 bottom) for 3hr and 96hr shows rapid paclitaxel-induced mitochondria vacuolization (red arrows) and membrane disruptions (yellow arrows), persisting up to 96hr, similar to paclitaxel+MMP-13 inhibitor. **(K)** Axonal HyPer oxidation is elevated after DMSO treatment for 3hr but not after prolonged treatment with either DMSO or paclitaxel, n=5-7 animals. *Abbreviations: B*=*basal keratinocyte*, *P*=*periderm*, *M*=*mitochondrion*, *Ax*=*axon*, *Ac*=*Actinotrichia*

Unlike in keratinocytes, TEM analysis of somatosensory axons within the epidermis showed that mitochondria continue to have distinct outer and inner membranes after treatment with either DMSO vehicle, paclitaxel or paclitaxel+DB04760, or CL-82198 for 3hr and 96hr (**Figure 3J**). Since many axons degenerated after 96hr treatment with paclitaxel, we assume that the remaining axons at this treatment phase are less sensitive to paclitaxel or damage induced in the skin. One possible explanation is that intact axons remain in areas of the skin where the matrix is less degraded. This would be consistent with our finding that only certain patches of basement membrane are degraded while others are still intact. One remarkable finding that our TEM analysis revealed is that, similar to mammalian DRG axons, paclitaxel treatment induced mitochondria vacuolization even after only 3hr treatment. These vacuoles persisted also in mitochondria of axons still present at 96hr of treatment, and in axons of fish in which either MMP-13 inhibitor (DB004760 or CL-82198) was co-administered. Since axon degeneration is prevented in the presence of MMP-13 inhibitors but vacuolization still occurs, it suggests that axonal mitochondrial vacuolization is not an underlying cause of axon degeneration. Moreover, the vacuoles appear rapidly and do not correlate with the onset of axon degeneration that occurs ~3-4 days later, which is substantial given that zebrafish develop from a single embryo into a hatched larva with most organs differentiated by 2dpf. In addition, vacuoles did not appear in keratinocyte mitochondria despite the fact that keratinocytes upregulate MMP-13 and induce axon degeneration. This finding indicates that vacuolization of mitochondria is not a cause of paclitaxel-induced axon degeneration.

Vacuolization has been suggested to induce mitochondrial damage and a disrupted mitochondrial permeability transmission pore in the inner membrane leading to reduced ATP production and mtROS leakage (19). Vacuolization has however also been observed in brown fat and liver tissue of starved rats (20), making it questionable whether these phenotypes are leading to cellular damage. If so, we would expect increased HyPer oxidation in axons. We assessed this by expressing HyPer-cyto under the CREST3 promoter in somatosensory neurons that innervate the epidermis **(Figure 3K)**. To our surprise, we could not detect increased H_2_O_2_ levels in epidermal sensory axons following paclitaxel treatment for 3hr, 48hr and 96hr. Interestingly, DMSO vehicle did initially, but transiently, increase HyPer oxidation. This effect was however undetectable after prolonged treatment. These data indicate that paclitaxel does not elevate axonal cytoplasmic H_2_O_2_ levels. The DMSO-dependent elevation of H_2_O_2_ might be due to increased membrane permeabilization, which could potentially be countered by a paclitaxel-mediated increase in axonal cytoskeletal (microtubule) rigidity.

Because of the similarities to mammalian paclitaxel-induced phenotypes in zebrafish, we wanted to test whether MMP-13-dependent axon degeneration is conserved in mammals. To induce PIPN, we injected rats and mice on alternating days with 2mg/kg and 4mg/kg paclitaxel in Cremophor/NaCl, respectively, over the course of 7 days (cumulative doses of 8mg/kg and 16mg/kg). Some animals received Cremophor/NaCl as vehicle control, or Cremophor/NaCl+CL-82198. As other studies have shown (9, 21-23), we were able to induce mechanical, thermal and cold nociceptive sensitivity in the paclitaxel-injected animals (**Figure 4A-D**). These phenotypes were prevented in paclitaxel-injected animals in which CL-82198 was co-administered. CL-82198/vehicle-injections produced comparable results to those injected with the vehicle alone. We also analyzed axon degeneration in the dorsal horn of injected rats using CGRP antibody staining, which showed reduced expression upon paclitaxel injections, and this effect was prevented in either vehicle or paclitaxel+CL-82198 injected animals (**Figure S6**).

**Figure 4.**
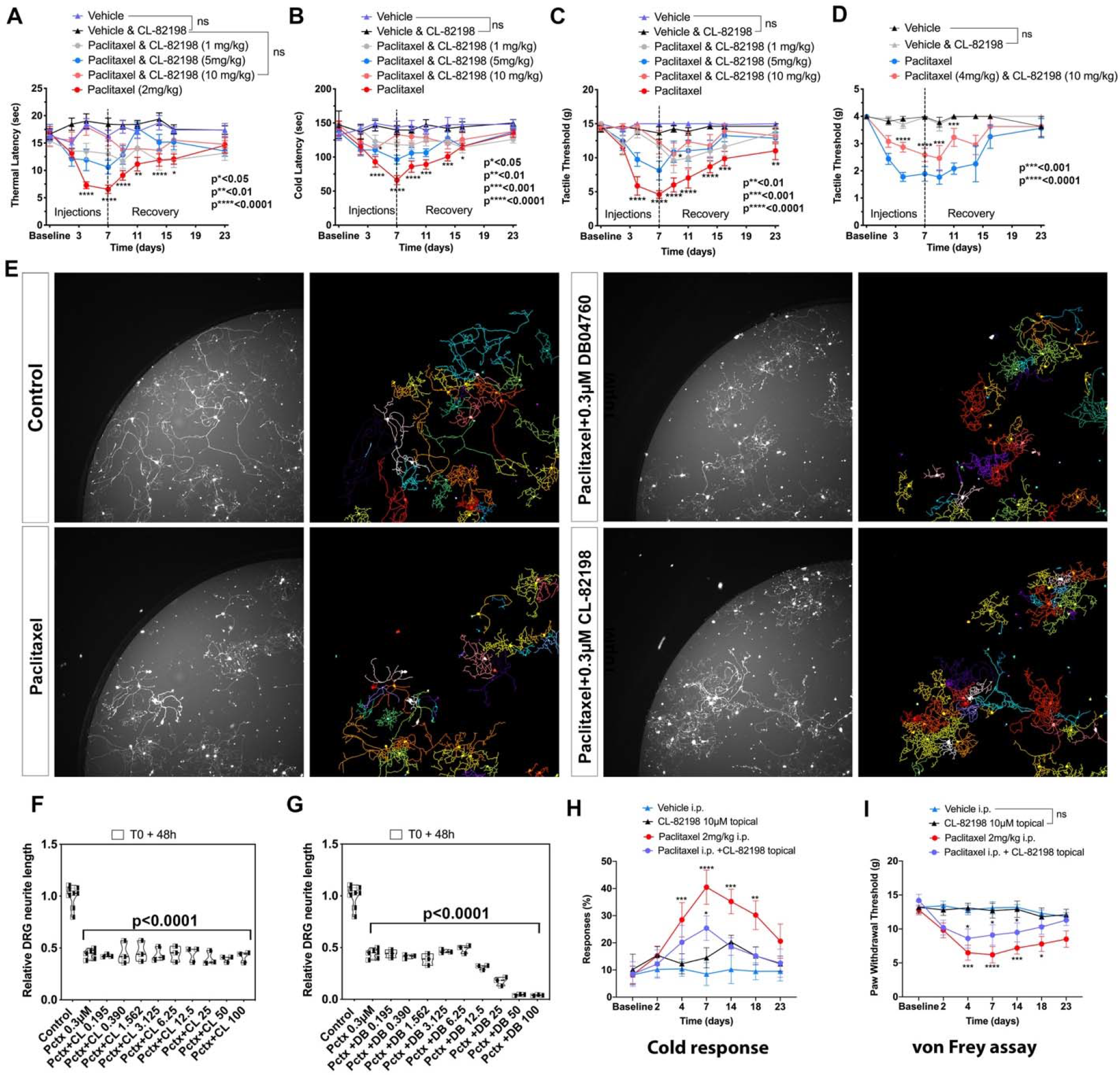
MMP-13 prevents paclitaxel-induced PIPN in rodents by neuron-extrinsic mechanism. **(A-D)** Increased sensitivity to heat (**A**), cold (**B**) and touch (**C, D**) following paclitaxel treatment of rats (**A-C**) and mice (**D**) compared with controls and upon injection of paclitaxel+CL-82198, n=6-14 animals. **(E-G)** DRG neurite tracings (**E**) and length quantification (**F, G**) shows decreased length upon treatment with either paclitaxel, paclitaxel+CL-82198, or paclitaxel+DB04760, n=3 biological replicates. (**H**, **I**) Increased cold response and tactile sensitivity is alleviated upon topical application of CL-82198 to the hind paw of paclitaxel-injected rats (n=3).

We previously showed in zebrafish that MMP-13 expression induced by paclitaxel was restricted to the epidermis and was not found in epidermal axons of sensory neurons (5). To determine whether MMP-13 dependence was neuron-intrinsic in mammals, we cultured mouse DRG neurons and examined neurite degeneration following treatment with DMSO vehicle, paclitaxel, paclitaxel+CL-82198, or paclitaxel+DB04760 at various concentrations. While paclitaxel induced neurite degeneration in the cultures, CL-82198 co-administration did not prevent degeneration, suggesting that similar to our zebrafish observations, MMP-13 is not functioning neuron-intrinsically (**Figure 4E-G**). Consistent with a skin-specific role for MMP-13, we found that topical application of 10μM CL-82198 to the hind paw of rats injected with paclitaxel led to a faster recovery from allodynia and cold sensitivity (**Figure 4H, I**). Thus, these findings indicate that MMP-13-dependent epidermal activity is conserved in both zebrafish and mammals, and likely also contributes to PIPN in human patients.

## Discussion

Our data suggests a model by which epidermal mitochondrial damage induces H_2_O_2_ production leading to increased MMP-13-dependent matrix degradation and axon degeneration (**Figure S7**). Mitochondria in the epidermis, and also in sensory axons, appear to be elongated upon paclitaxel treatment, likely related to defects in the fusion or fission machinery. Previous studies showed that H_2_O_2_ promotes mitochondrial fission, but our data points to increased filamentation, arguing against a model in which H_2_O_2_ damages mitochondria. Instead, mitochondrial damage appears to cause H_2_O_2_/ROS formation leading to downstream effects, such as MMP-13 activation. Strikingly, although mitochondria damage similarly occurs in axons, we don’t believe this to be the cause of axon degeneration. We showed that pharmacological MMP-13 inhibition can prevent axon degeneration (5), yet, MMP-13 inhibition does not prevent axonal mitochondrial vacuolization and morphological changes. It is also interesting that axons accumulate elongated mitochondria but at the same time significantly increase the number of mitochondria per branch segment. It is possible that axons have an increased ability to undergo fusion and fission at an accelerated rate as a means to increase mitochondrial health, whereas this ability is inhibited in keratinocytes.

Why do epidermal mitochondria contribute to MMP-13 expression, but sensory neurons do not upregulate MMP-13? The simplest explanation is that regulatory elements leading to MMP-13 expression are not present in these neurons, and thus axons cannot upregulate MMP-13. This is consistent with our mouse DRG *in vitro* culture data (**Figure 4E-G**). However, it has been well established that H_2_O_2_/ROS can upregulate other MMPs, which could in principle contribute to axon degeneration. But because H_2_O_2_/ROS levels are low, as determined in our HyPer experiments (**Figure 3K**), MMPs won’t be activated in the context of paclitaxel treatment. Another pertinent question is why axons have low H_2_O_2_/ROS levels while keratinocytes do not. Perhaps axons have the ability to better regulate H_2_O_2_/ROS through means of antioxidant complexes that scavenge H_2_O_2_/ROS. Evidence for this theory stems from the fact that axons can survive and even regenerate upon exposure to injury-released H_2_O_2_/ROS from keratinocytes (14) whereas keratinocytes that are exposed to prolonged H_2_O_2_/ROS levels might be more vulnerable to damage. Thus, epidermis-innervating somatosensory neurons must have evolved a better mechanism to tolerate long-term exposure to H_2_O_2_/ROS levels. This difference could hinge on the H_2_O_2_-dependent expression of MMP-13. Under conditions of paclitaxel treatment, keratinocytes utilize the same mechanisms of MMP-13 upregulation as after skin injury, where MMP-13 is localized to the leading edge of migrating keratinocytes during the re-epithelialization step of the wound (24). Because MMP-13 actions are transient, this process is coordinated with the healing of the wound, after which H_2_O_2_/ROS and MMP-13 are downregulated. However, if MMP-13 remains high, continuous matrix degradation under homeostatic conditions will damage epidermal cells and ultimately the axons.

How is mitochondrial damage induced by paclitaxel in keratinocytes and axons? Previous studies demonstrated that paclitaxel can directly damage isolated mitochondria of neuroblastoma cells (8) and liver cells (25), and thus it could also exert direct effects on mitochondria in cell types, such as keratinocytes and axons. This also indicates that microtubule damage, as observed in cells treated with paclitaxel, may not contribute significantly to axon degeneration. While microtubule damage is a desired effect during cancer treatment, neurons do not divide, and keratinocytes in zebrafish do rarely divide during larval stages. However, basal keratinocytes divide frequently in the mammalian and the adult zebrafish epidermis, and paclitaxel treatment may have additional effects in dividing epithelia that could lead to further damage, though this remains to be shown. Nevertheless, our mammalian data and previous adult zebrafish data (5) showing the reversal of behavioral symptoms and axon degeneration induced by paclitaxel in the presence of MMP-13 inhibitor suggests that MMP-13 is a primary inducer of axon degeneration.

## Supporting information

Cirrincione et al. Supplementary data file

## Acknowledgments

We thank Pete Finger at The Jackson Laboratory for his assistance with the TEM studies. We also thank the fish facility staff at MDI Biological Laboratory and the University of Miami for their excellent zebrafish care.

