## Supplementary material for "Paclitaxel-induced peripheral neuropathy is caused by epidermal ROS and mitochondrial damage through conserved MMP-13 activation": Cirrincione et al. Supplementary data file

**Supplementary file**

**Supplementary Figure legends**

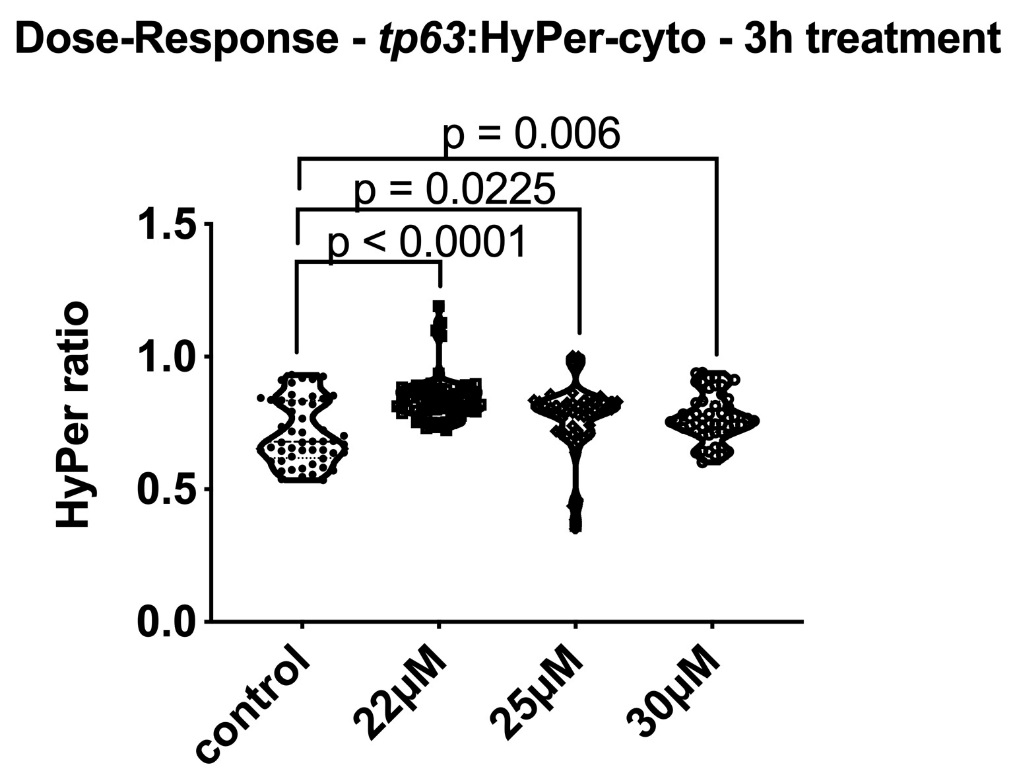

**Figure S1. Dose-response curve of paclitaxel-induced H_2_O_2_ formation in basal keratinocytes.** Zebrafish larvae at 2dpf were treated for 3hr with either vehicle or 22, 25, and 30µM paclitaxel, with maximal difference in H_2_O_2_ levels, when compared to vehicle controls, at 22µM, n=5-7 animals.

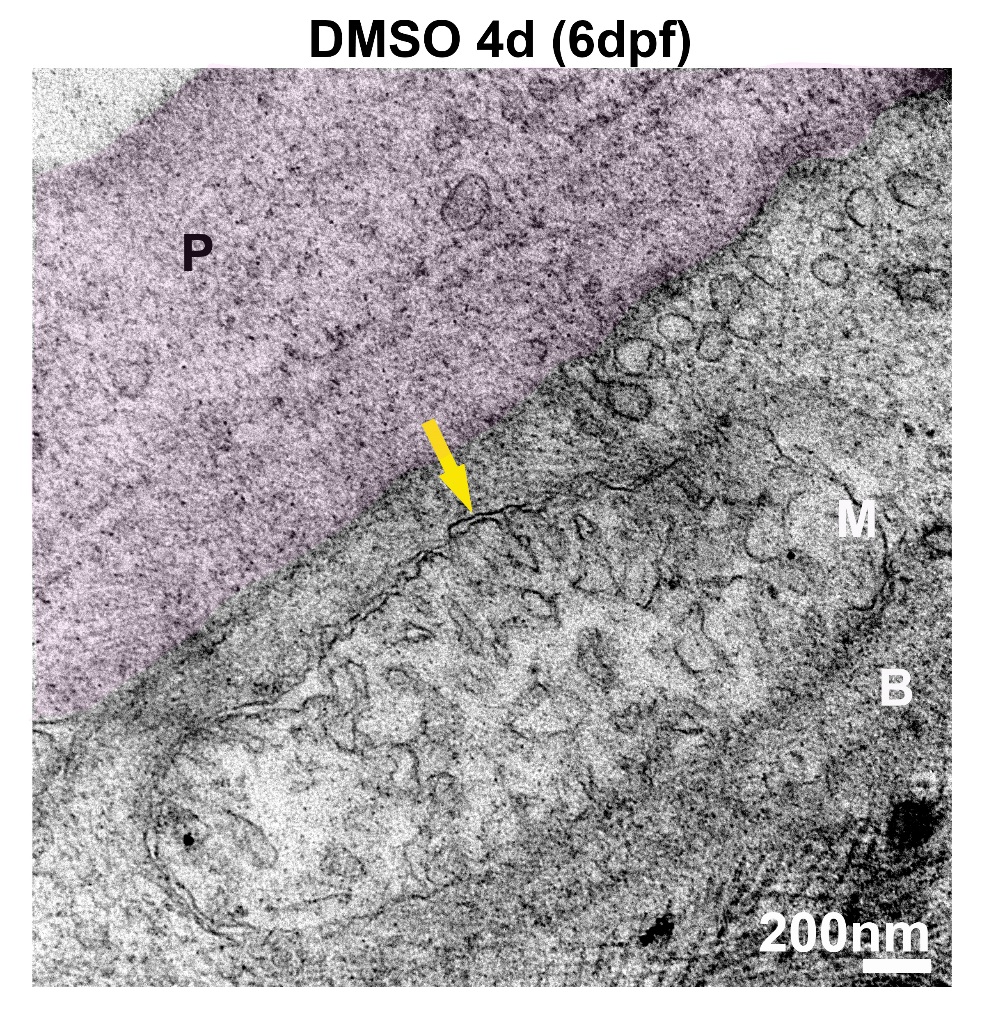

**Figure S2.** **Transmission electron microscopy showing a mitochondrion in a basal keratinocyte following vehicle treatment for 96hr**. Zebrafish larva treated with 0.09% DMSO vehicle for 4d (until 6dpf). The outer and inner mitochondrial membrane (yellow arrow) and cristae are clearly visible. The periderm is shaded. *Abbreviations: P=periderm, M=mitochondrion, B=basal cell.*

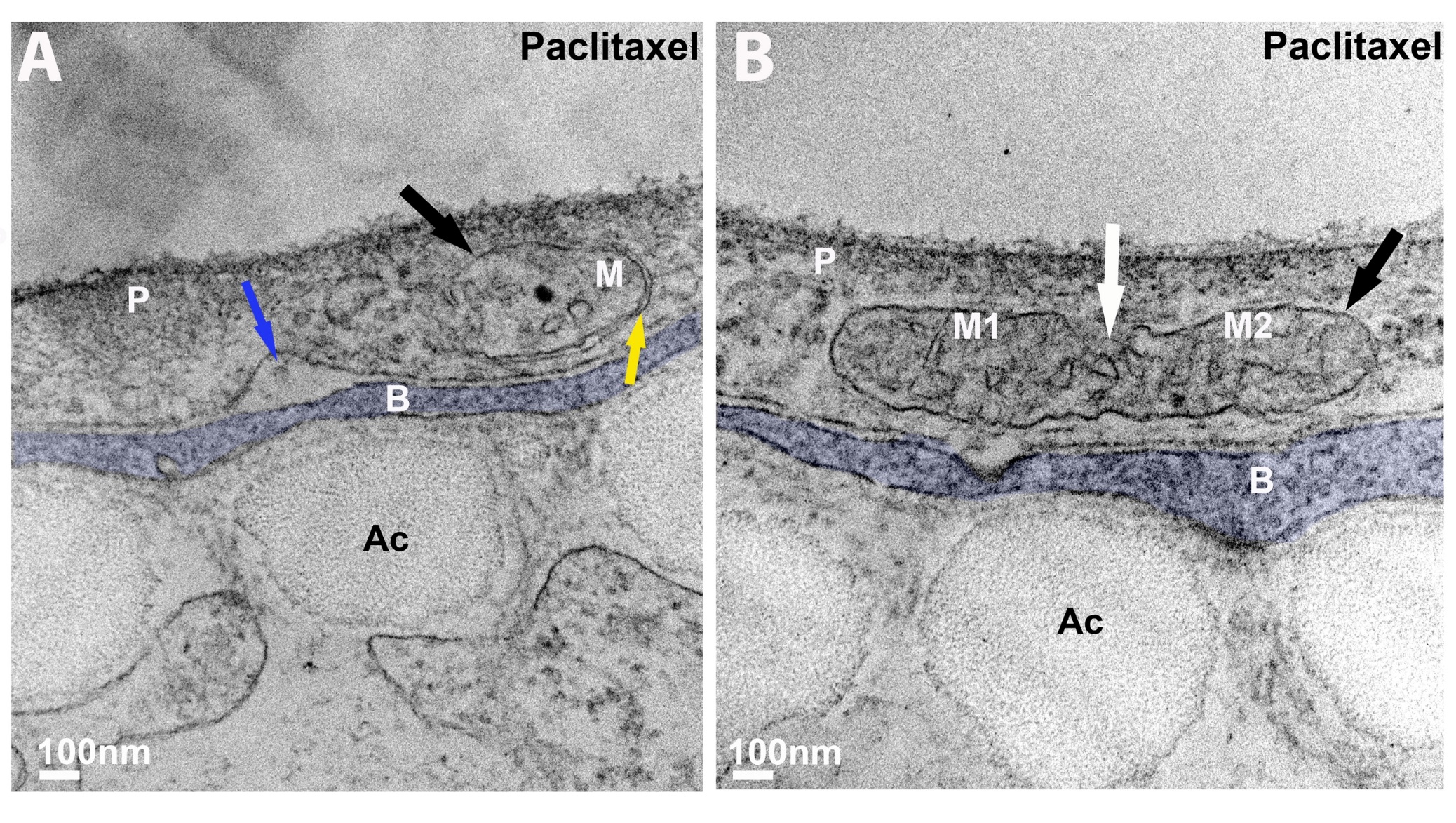

**Figure S3. Transmission electron microscopy showing a mitochondrion in basal keratinocytes following paclitaxel treatment for 96hr**. (**A, B**) Zebrafish larvae treated with 23µM paclitaxel for 4d (until 6dpf). The outer and inner membrane are visible in some parts of the mitochondrion (yellow arrow), and absent in others (black arrow**, B**). The cristae are visible in (**B**) and barely visible in (**A**). Electron-dense puncta are present, which are not observed in controls (**Figure S3**) or following co-administration of DB04760 or CL-82198 (**Figure S5**). The mitochondrion in (**B**) appears to be fused to another one (white arrow). A large gap where normally an axon would reside is apparent in (**A,** blue arrow). The basal cells are shaded. *Abbreviations: P=periderm, M=mitochondrion, B=basal cell, Ac=Actinotrichia.*

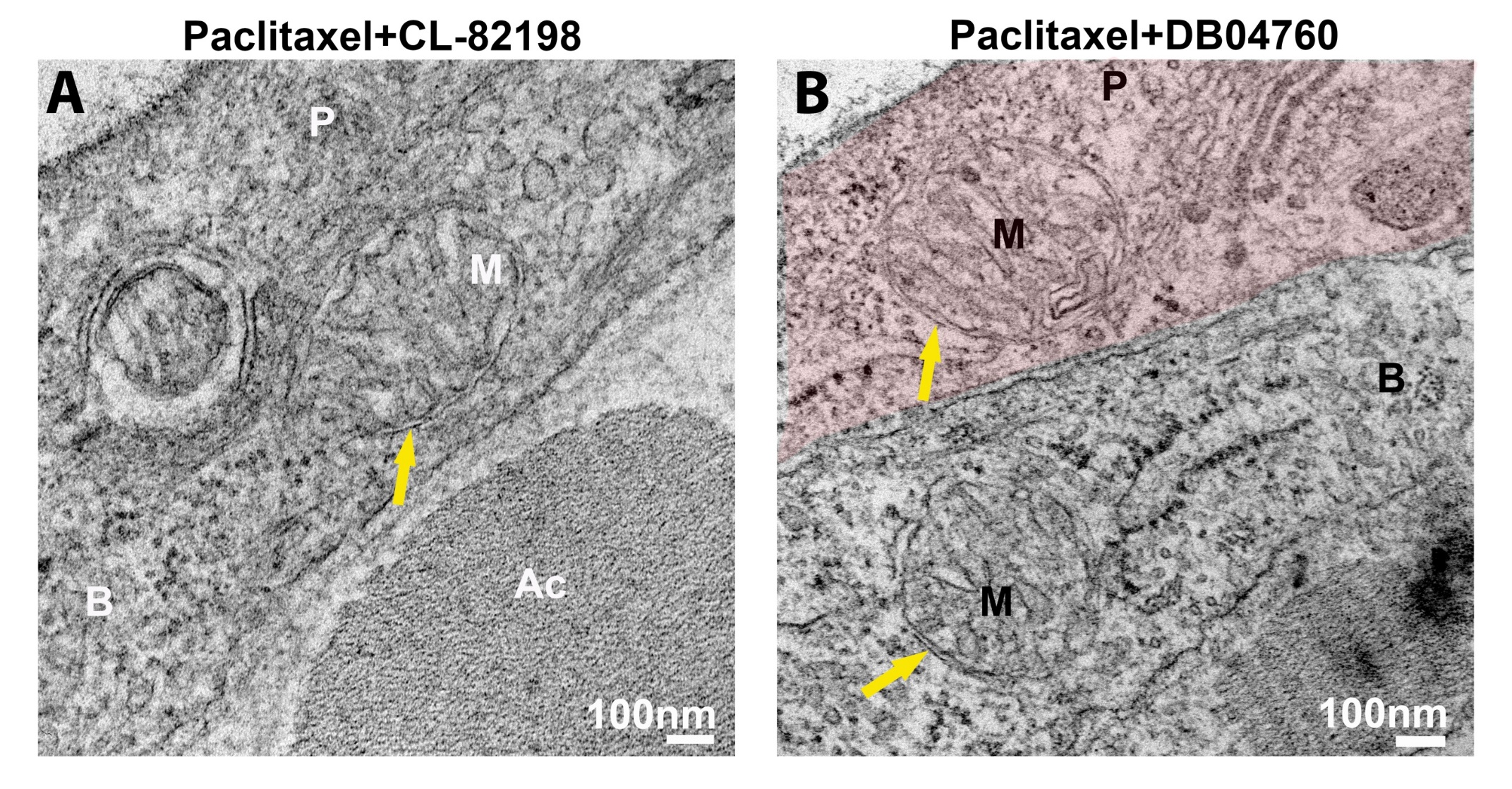

**Figure S4. Transmission electron microscopy showing a mitochondrion in basal keratinocytes following treatment with paclitaxel and CL-82198 or DB04760 for 96hr**. Zebrafish larva treated with 23µM paclitaxel and either 10µM CL-82198 (**A**) or DB04760 (**B**) for 4d (until 6dpf). The outer and inner mitochondrial membranes and cristae are clearly visible (yellow arrows) when MMP-13 was inhibited. The periderm is shaded. *Abbreviations: P=periderm, M=mitochondrion, B=basal cell, Ac=Actinotrichia.*

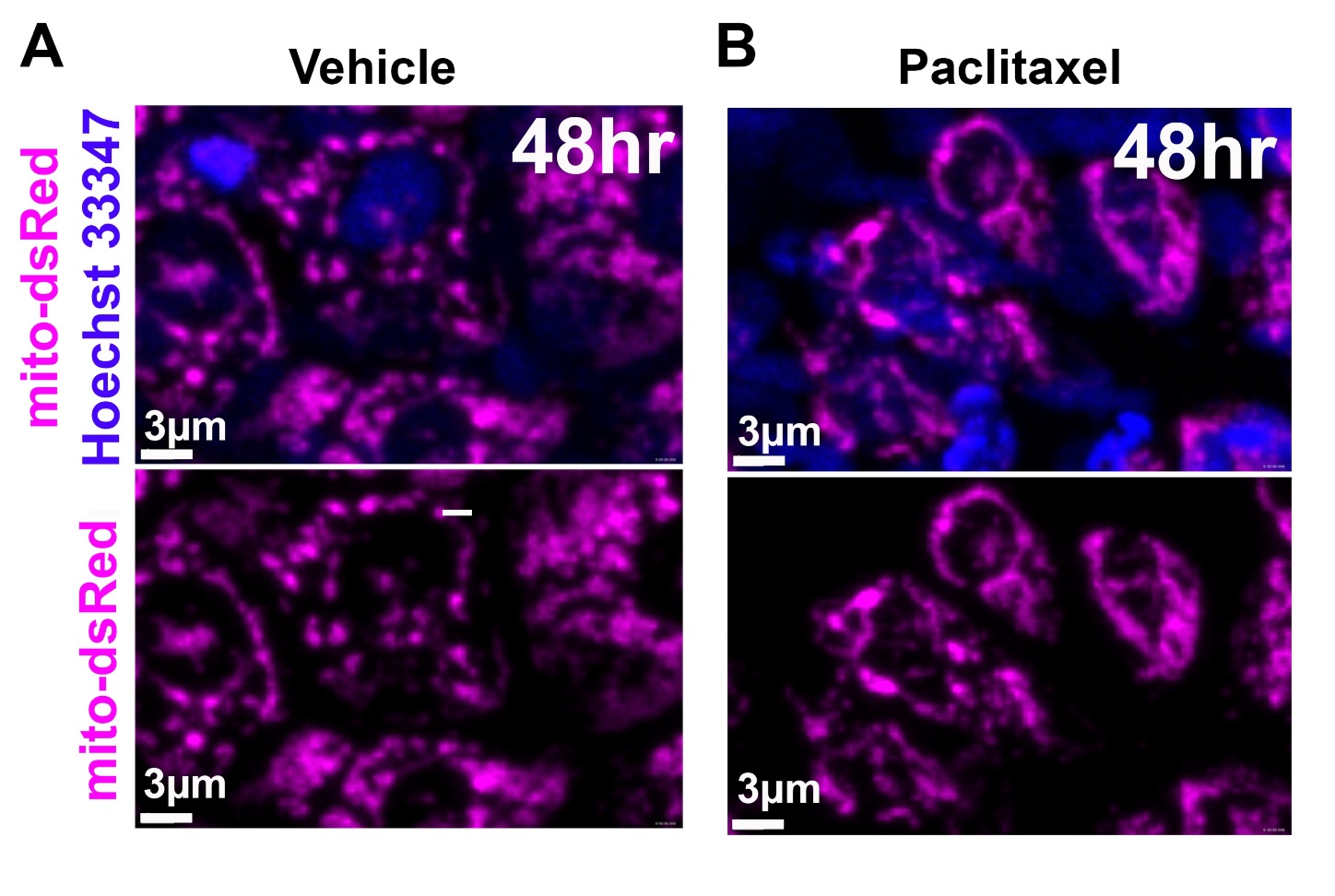

**Figure S5. Aberrant morphology of keratinocyte mitochondria following paclitaxel treatment for 48hr starting at 2dpf. (A, B)** Transgenic zebrafish UAS-mito-dsRed was injected at the 1-cell stage with *krt4*:Galv4VP16 to label mitochondria. Filamentous mitochondria are visible following paclitaxel treatment (B) but not upon vehicle treatment (A).

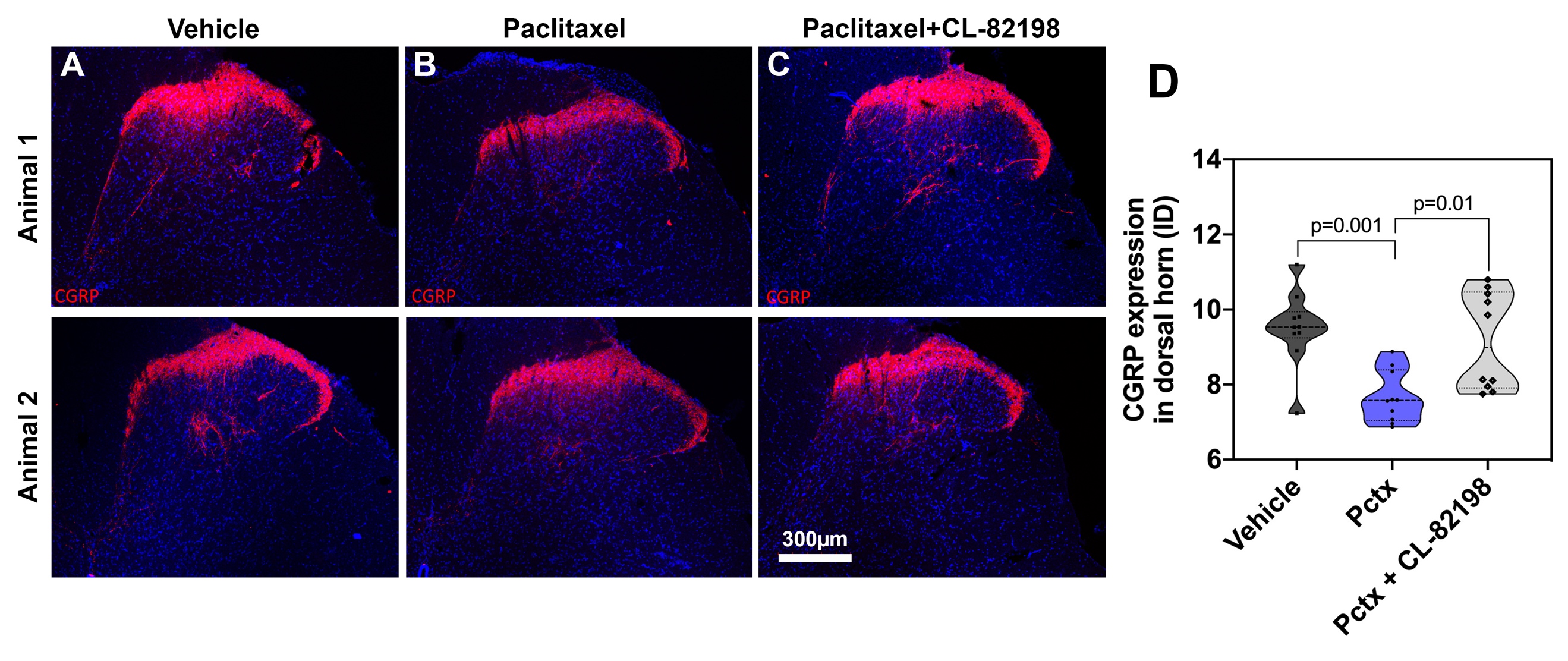

**Figure S6. CL-82198 prevents paclitaxel-induced axon degeneration upon i.p. injection and topical application to the epidermis of Sprague-Dawley rats. (A-C)** CGRP staining to label the dorsal horn (red) overlaid with DAPI nuclear staining shows reduced staining upon paclitaxel treatment. (**D**) Quantification of fluorescence confirms the observations in **A-C**, n=5 animals.

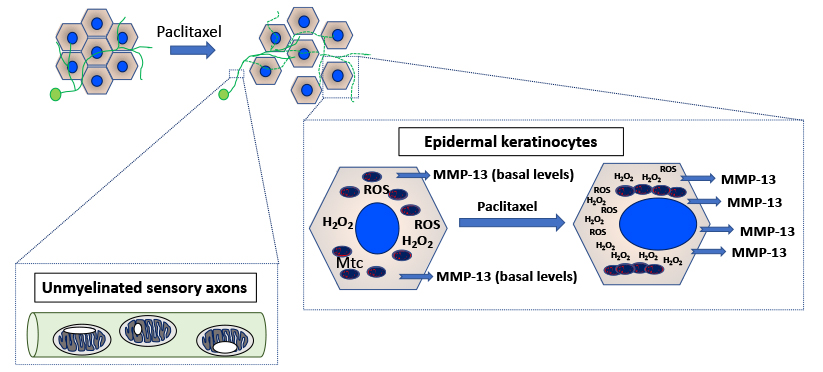

**Figure S7. Model for paclitaxel-induced peripheral neuropathy via mitochondrial ROS formation in the epidermis.** Paclitaxel rapidly damages mitochondria in epidermal keratinocytes, which leads to increased ROS/H_2_O_2_ formation and upregulation of MMP-13. MMP-13 dependent matrix degradation in certain stretches of the basement membrane affects unmyelinated axons that traverse the basement membrane into the epidermis, leading to axonal degeneration. Axonal mitochondria are also rapidly changed in morphology, such as the appearance of vacuoles and increased size upon paclitaxel treatment. However, these mitochondria do not appear to contribute to H_2_O_2_ leakage into the axonal cytoplasm, and MMP-13 has no role in axon degeneration.
